# Linkage disequilibrium dependent architecture of human complex traits reveals action of negative selection

**DOI:** 10.1101/082024

**Authors:** Steven Gazal, Hilary K. Finucane, Nicholas A Furlotte, Po-Ru Loh, Pier Francesco Palamara, Xuanyao Liu, Armin Schoech, Brendan Bulik-Sullivan, Benjamin M Neale, Alexander Gusev, Alkes L. Price

## Abstract

Recent work has hinted at the linkage disequilibrium (LD) dependent architecture of human complex traits, where SNPs with low levels of LD (LLD) have larger per-SNP heritability after conditioning on their minor allele frequency (MAF). However, this has not been formally assessed, quantified or biologically interpreted. Here, we analyzed summary statistics from 56 complex diseases and traits (average N = 101,401) by extending stratified LD score regression to continuous annotations. We determined that SNPs with low LLD have significantly larger per-SNP heritability. Roughly half of the LLD signal can be explained by functional annotations that are negatively correlated with LLD, such as DNase I hypersensitivity sites (DHS). The remaining signal is largely driven by our finding that common variants that are more recent tend to have lower LLD and to explain more heritability (*P* = 2.38 × 10^−104^); the youngest 20% of common SNPs explain 3.9x more heritability than the oldest 20%, consistent with the action of negative selection. We also inferred jointly significant effects of other LD-related annotations and confirmed via forward simulations that these annotations jointly predict deleterious effects. Our results are consistent with the action of negative selection on deleterious variants that affect complex traits, complementing efforts to learn about negative selection by analyzing much smaller rare variant data sets.

## Introduction

Estimating the heritability explained by SNPs^1,2^, and its distribution across chromosomes^3,4^, allele frequencies^5^ and functional regions^6–10^, has yielded rich insights into the polygenic architecture of human complex traits. Recent work has hinted at linkage disequilibrium (LD) dependent architectures, defined as a dependence of causal effect sizes on levels of LD (LLD) after conditioning on minor allele frequency (MAF), for several complex traits. LD-dependent architectures bias SNP-heritability estimates^11^, and downward biases have been observed for several traits^11–13^, suggesting larger causal effect sizes for genetic variants with low LLD. Indeed, heritability is enriched in functional annotations such as DNase I hypersensitivity sites (DHS)^7^, histone marks^8,10^, and regions with high GC-content^9^, which all have low LLD^7,14,15^. On the other hand, regions of low recombination rate, which have high LLD, are enriched for exonic deleterious and disease-associated variants^16^, suggesting an LD-dependent architecture of opposite effect.

Despite these observations, LD-dependent architectures have not been formally assessed, quantified, or biologically interpreted. Understanding which biological processes shaping the LD patterns of the genome are most directly linked to complex traits is challenging, as many of the corresponding annotations are correlated with each other. To investigate LD-dependent architectures, we extended stratified LD score regression^8^, a method that partitions the heritability of a set of binary genomic annotations using GWAS summary statistics, to continuous-valued annotations; our method produces robust results in simulations. We applied our method to a broad set of LD-related annotations, including LLD, predicted allele age and recombination rate, to analyze summary statistics from 56 complex traits and diseases (average *N* = 101,401), including 18 traits from the 23andMe, Inc. research database and 15 traits from the UK Biobank. We inferred jointly significant effects of several LD-related annotations on per- SNP heritability, including predicted allele age: common variants that are more recent tend to have lower LLD and to explain more heritability, which is consistent with the action of negative selection since selection has had less time to eliminate recent weakly deleterious variants. We confirmed via forward simulations that allele age, as well as other LD-related annotations associated to per-SNP heritability, jointly predict the deleterious effects of a variant. Our results implicate the action of negative selection on deleterious variants that affect complex traits.

## Results

### Overview of methods

Stratified LD score regression^8^ is a method for partitioning heritability across overlapping binary annotations using GWAS summary statistics. The idea of this method is that, for a polygenic trait, LD to an annotation that is enriched for heritability will increase the *χ^2^* statistic of a SNP more than LD to an annotation that is not enriched for heritability. We extended stratified LD score regression to quantify effects on heritability of continuous-valued (and/or binary) annotations. Here, the idea is that if a continuous annotation *a* is associated to increased heritability, LD to SNPs with large values of *a* will increase the *χ^2^* statistic of a SNP more than LD to SNPs with small values of *a*.

More precisely, the expected *χ^2^* statistic of SNP *j* can be written as

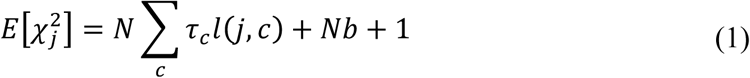
where 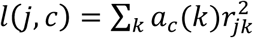 is the LD score of SNP *j* with respect to continuous values *a_c_* (*k*) of annotation *a_c_*, *r_jk_* is the correlation between SNP *j* and *k* in a reference panel (e.g. Europeans from 1000 Genomes^17^), *N* is the sample size of the GWAS study, *τ_c_* is the effect size of annotation *a_c_* on per-SNP heritability (conditioned on all other annotations), and *b* is a term that measures the contribution of confounding biases^18^. We standardize estimated effect sizes 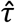 to report per-standardized-annotation effect sizes *τ*\*, defined as the proportionate change in per-SNP heritability associated to a 1 standard deviation increase in the value of the annotation; we note that *τ\** can be compared across annotations and across traits. Analogous to ref. ^8^, standard errors on estimates of *τ\** are computed using a block jackknife (see Online Methods). We have released open-source software implementing the method (see URLs).

We applied our extension of stratified LD score regression to LLD annotations, MAF-adjusted via MAF-stratified quantile-normalized LD score, as well as other LD-related annotations including predicted allele age and recombination rate; we included 10 MAF bins as additional annotations in all analyses to model MAF-dependent architectures. We also considered functional annotations from a “baseline model”^8,19^ including 28 main annotations such as coding, conserved, DHS and histone marks (59 total annotations; see Online Methods).

Although stratified LD score regression has previously been shown to produce robust results using binary annotations^8^, we performed additional simulations to confirm that our extension of stratified LD score regression produces robust results using continuous-valued LD-related annotations, and specifically that analyzing LD-related annotations using an LD-based method is appropriate (see Online Methods).

### SNPs with low LLD have larger per-SNP heritability

We applied our extension of stratified LD score regression to GWAS summary statistics from 56 complex traits and diseases, including 18 traits from 23andMe and 15 traits from UK Biobank (average *N* = 101,401); for five traits we analyzed multiple data sets, leading to a total of 62 data sets analyzed (Table S1). The standardized effect sizes *τ\** for the LLD annotation were consistently negative in all 62 data sets analyzed (Figure 1 and Table S2). In a meta-analysis across 31 independent traits, excluding genetically correlated traits^20^ in overlapping samples (Table S3; average *N* = 84,686, see Online Methods), the LLD annotation was highly statistically significant (*τ\** = -0.30, s.e. = 0.02; *P* = 2.42 × 10^−80^), confirming that SNPs with low MAF-adjusted level of LD have larger per-SNP heritability. We also investigated two alternative MAF-adjusted measures of level of LD, using a sliding window approach to quantify the level of LD in a genomic region (LLD-REG)^13^ and using the D’ coefficient instead of the squared correlation to compute LD scores (LLD-D’); we observed smaller but still significant effects for LLD-REG (*τ\** = -0.22, s.e. = 0.02; *P* = 2.86 × 10^−44^) and LLD-D’ (*τ\** = -0.15, s.e. = 0.02; *P* = 2.22 × 10^−12^).

**Figure 1:**
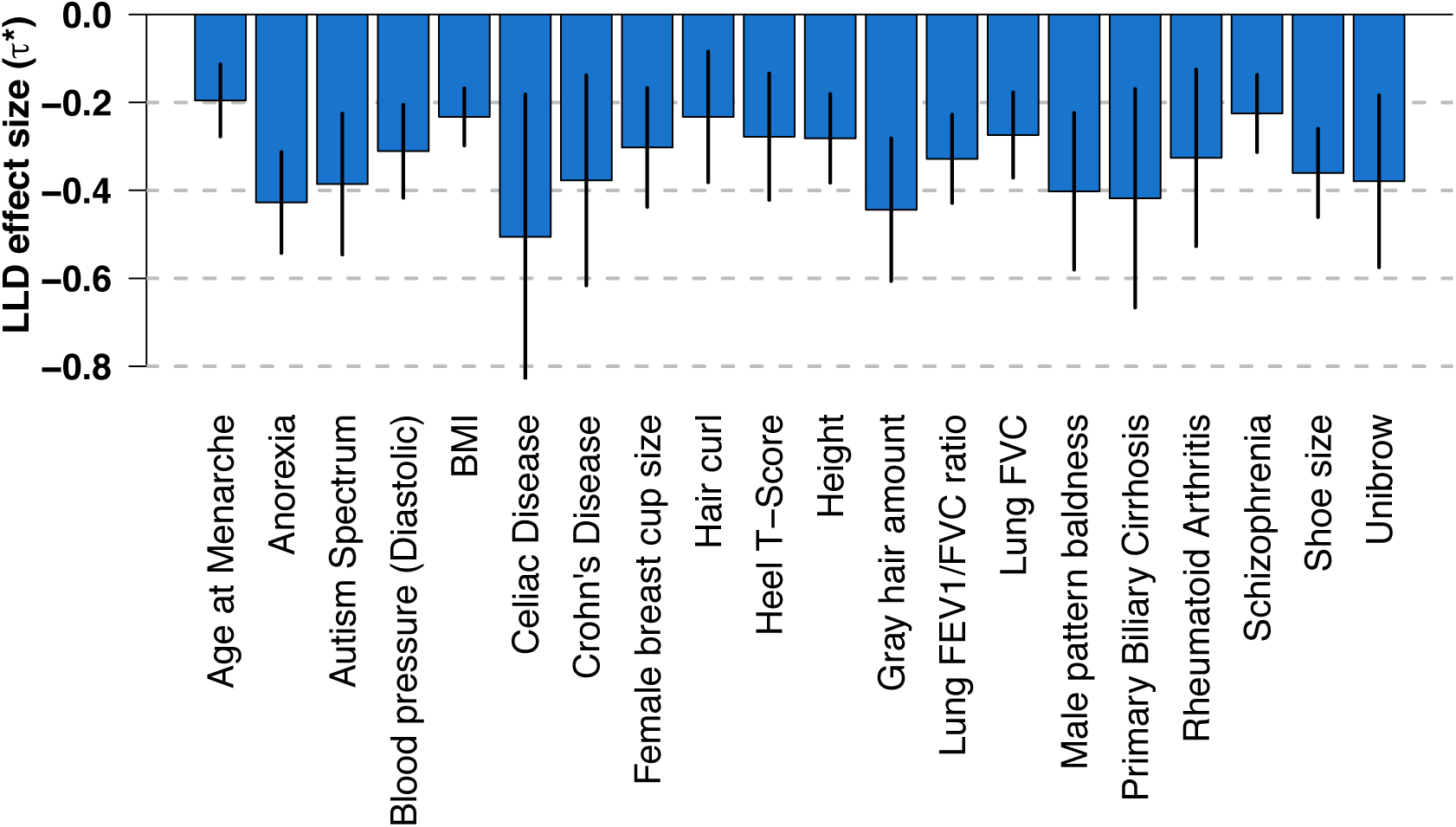
Effect size of MAF-adjusted level of LD (LLD) on 20 highly heritable complex traits. Results are displayed for 20 traits with the highest SNP-heritability (subject to low genetic correlation^20^ between traits). Numerical results for all 56 complex traits are reported in Table S2. Error bars represent jackknife 95% confidence intervals.

### Correlations between LLD and other LD-related annotations

We investigated other LD-related annotations including MAF-adjusted allele age as predicted using ARGweaver^21^, MAF-adjusted LLD measured in African populations (LLD-AFR), recombination rate^22,23^, nucleotide diversity^15^, a background selection statistic (McVicker B-statistic)^24^, GC-content^15^, CpG dinucleotide content, replication timing^25^, centromeres and telomeres^15^. We used the Oxford recombination map^23^ and a window size of ±10kb for recombination rate, and window sizes of ±10kb for nucleotide diversity, ±1Mb for GC-content, ±50kb for CpG-content, ±5Mb for centromeres and first/last 10Mb for telomeres, as these choices produced the most significant signals, although other choices produced similar results (see below and Online Methods). We also considered the 28 main functional annotations from our baseline model˙. Many of these annotations are highly correlated with LLD and with each other (Figure 2 and Table S4); these correlation patterns inform the interpretation of our heritability results below. In particular, nearly all of the functional annotations from the baseline model are negatively correlated with LLD, with the strongest negative correlations (-0.20 < *r* < -0.10) for histone marks (H3K27ac, H3K4me1 and H3K9ac), conserved regions (GERP NS) and super enhancers; only repressed regions (*r* = 0.05; depleted for heritability^8^) and transcribed regions (*r* = 0.02) exhibit positive correlations.

**Figure 2:**
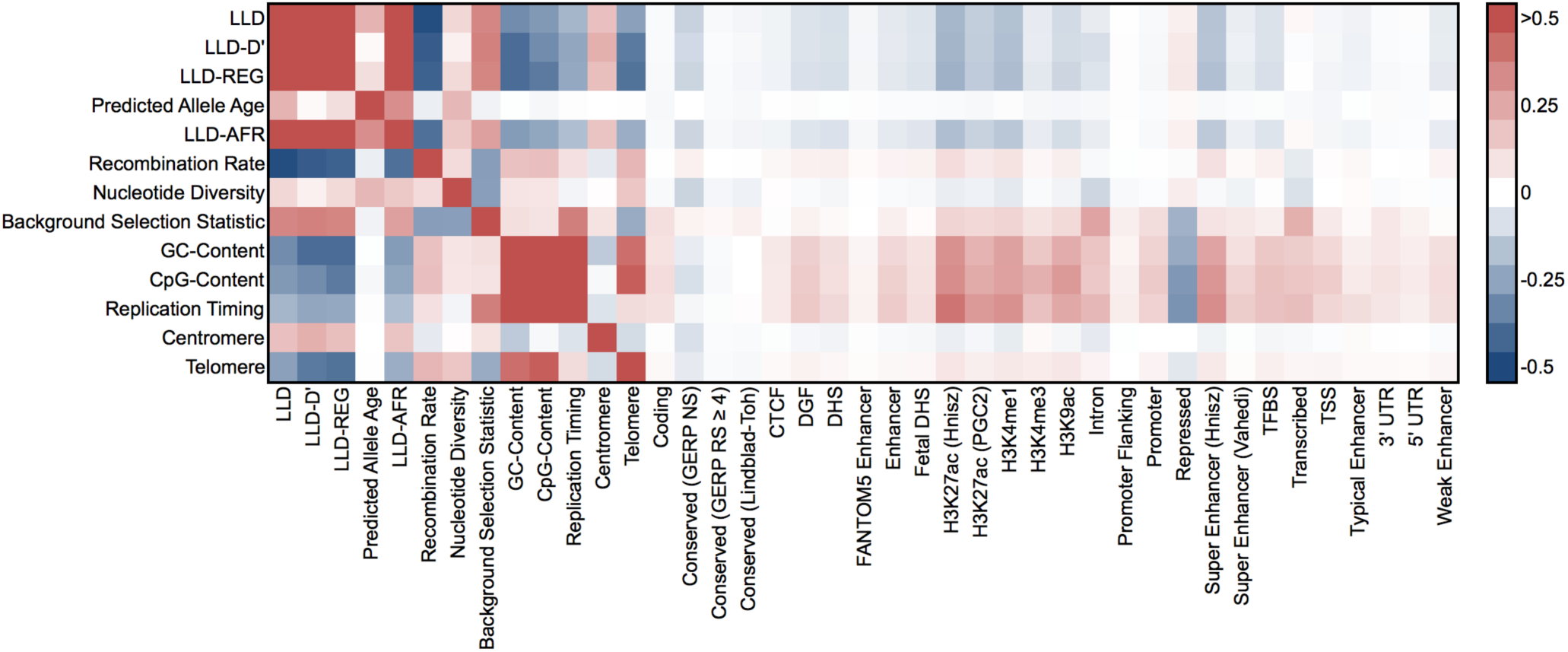
Correlations between LD-related and functional annotations. We report correlations computed on common SNPs (MAF ≥ 5%). LLD, LLD-D’, LLD-REG, predicted allele age and LLD-AFR annotations are MAF-adjusted. Numerical results are reported in Table S4.

One surprising observation was that predicted allele age was positively correlated with LLD (*r* = 0.22; more recent SNPs have lower LLD), whereas a negative correlation might be expected since the LD between two SNPs decays with time. To confirm this observation, we performed coalescent simulations^26^ using a realistic demographic model for African and European populations^27^ (see Online Methods). We observed that while the LLD of a SNP defined using a fixed set of older SNPs decreases with allele age, older SNPs acquire additional LD with more recent SNPs; the latter effect leads to a positive correlation between predicted allele age and LLD (Figure S1 and Figure S2). We also observed, in both real data and simulations, that allele age is more strongly correlated to LLD-AFR than LLD, as demographic events (e.g. bottlenecks) that occurred in European populations distort the relationship between LLD and allele age.

### Multiple LD-related annotations impact complex trait architectures

We applied our extension of stratified LD score regression to each of the 13 LD-related annotations defined above, analyzing each annotation in turn. We meta-analyzed the results across 31 independent traits (Figure 3a, Table S2 and Table S3). All annotations except telomeres were highly significant after correction for multiple testing (Table S3), and eight of the remaining 12 annotations remained significant when fitted jointly (Table S5 and Table S6). The predicted allele age (*τ\** = -0.78, s.e. = 0.03; *P* = 6.27 × 10^−175^) and nucleotide diversity (*τ\** = -0.78, s.e. = 0.04; *P* = 1.79 × 10^−79^) annotations produced the largest absolute standardized effect size. Interestingly, SNPs in high recombination rate regions (corresponding to low LLD; *r* = -0.49) have smaller per-SNP heritability (*τ\** = -0.54, s.e. = 0.06; *P* = 2.39 × 10^−18^), which is inconsistent with the direction of the LLD effect but consistent with the fact that negative selection is more effective in high recombination rate regions as a consequence of the Hill-Robertson effect^28^. Thus, per-SNP heritability is most enriched in SNPs with low LLD in low recombination rate regions, and the opposing effects of these two annotations are stronger when they are conditioned on each other (Figure S3). Opposing effects were also observed for the background selection statistic annotation, which is positively correlated to LLD (*r* = 0.35) but has the opposite direction of effect (*τ\** = 0.51, s.e. = 0.05; *P* = 5.06 × 10^−26^).

**Figure 3:**
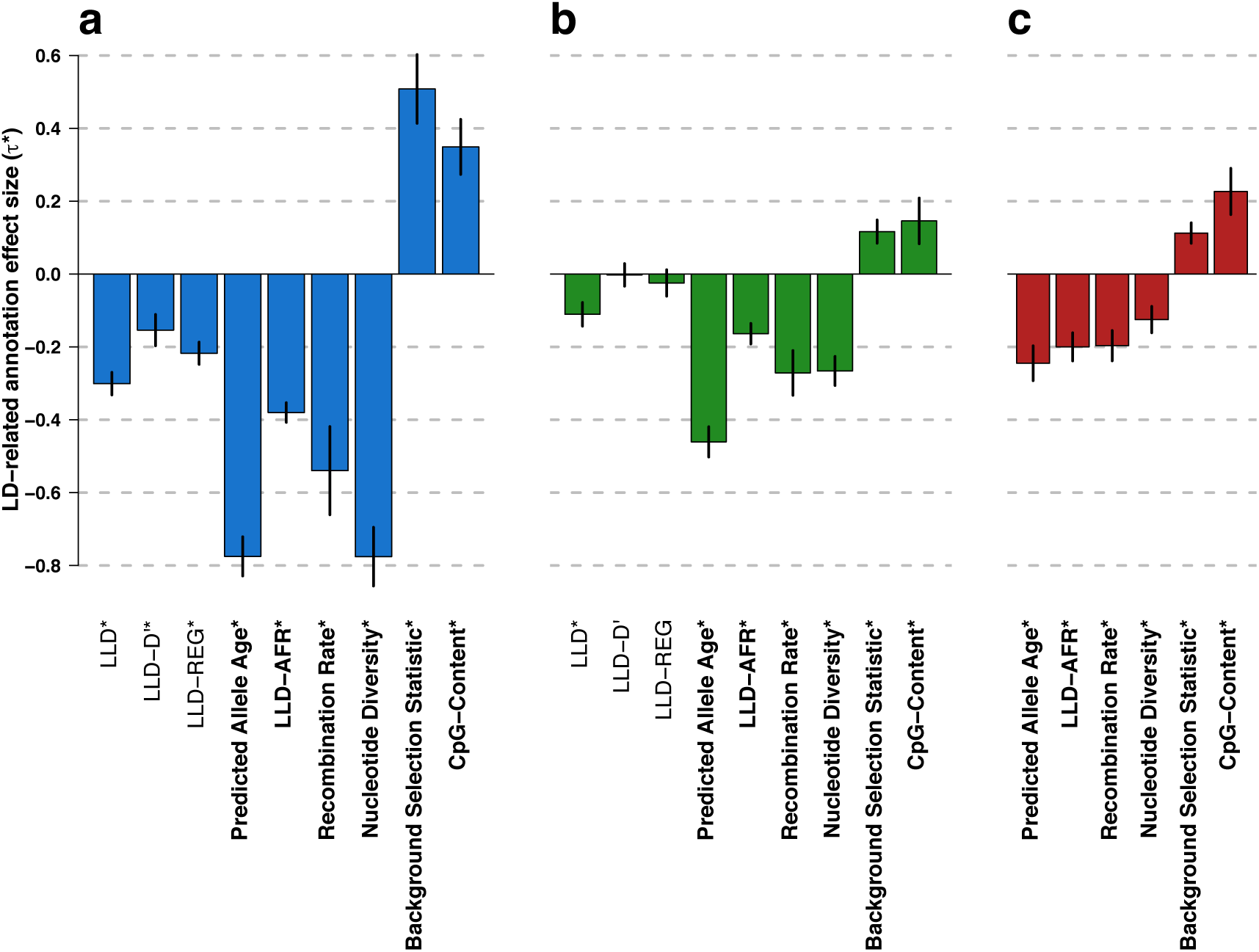
Effect size of LD-related annotations meta-analyzed over 31 independent traits. **(a)** Meta-analysis results for 9 LD-related annotations. **(b)** Meta-analysis results for nine LD-related annotations, conditioned on baseline model. **(c)** Meta-analysis results for six LD-related annotations conditioned on each other and on baseline model. Results are displayed for the six LD-related annotations that are jointly significant when conditioned on each other and on the baseline model (see (c)). In (a) and (b) only, results are also displayed for the remaining LLD annotations. Numerical results for all annotations analyzed are reported in Table S3 for (a) and (b), and Table S8 for (c). Numerical results for all 56 complex traits are reported in Table S2 for (a), Table S7 for (b), and Table S9 for (c). Asterisks indicate significance at *P* < 0.05 after Bonferroni correction (0.05/43, 0.05/43, and 0.05/6 for (a), (b), (c), respectively). Error bars represent 95% confidence intervals

In order to assess how much of the LLD effect is explained by known functional annotations (and because results of stratified LD score regression may be biased in the presence of unmodeled functional annotations^8^), we analyzed each of the 13 LD-related annotations while conditioning on the 59 functional annotations of the baseline model (Figure 3b, Table S3 and Table S7). The effect size of the LLD annotation remained highly significant but was smaller in magnitude (*τ\** = -0.11, s.e. = 0.02; *P* = 2.57 × 10^−11^), primarily due to its correlation with DHS (Figure S4). Thus, more than half of the initial LLD signal is explained by known functional annotations. The LLD-REG annotation^13^ was no longer significant in this analysis (*P* = 0.19), indicating that the regional LLD signal is entirely explained by known functional annotations. Predicted allele age produced the largest absolute standardized effect size and the most significant signal (*τ\** = -0.46, s.e. = 0.02; *P* = 2.38 × 10^−104^); the sign of this effect was consistent across 55 out of 56 traits (positive but not significantly different from zero for Hb1AC; Table S7). This indicates that more recent alleles have larger per-SNP heritability after conditioning on both MAF and known functional annotations. Many other LD-related annotations remained significant (after correction for multiple testing) in the conditional analysis, although LLD-D’, replication timing and centromeres were no longer significant (Table S3 and Figure S4).

Finally, we built a model consisting of the 59 functional annotations from the baseline model and the six LD-related annotations that remained significant (after correction for multiple testing) when conditioned on each other as well as the baseline model (Figure 3c, Table S8 and Table S9); we call this model the baseline-LD model (see Online Methods). We determined that this model produced similar results when using different window sizes for windows-based annotations (e.g. recombination rate, nucleotide diversity and CpG-content) or different data sources for recombination rate (Figure S5), when performing derived allele frequency (DAF) adjustment instead of MAF adjustment, when using UK10K^29^ (instead of 1000 Genomes) as the reference panel (Figure S6), and across different data sets for the same trait (Figure S7). Predicted allele age remains the annotation with the largest absolute standardized effect size (*τ\** = -0.24, s.e. = 0.02; *P* = 1.08 × 10^−23^), but its effect size decreased due to its high correlation with the LLD-AFR annotation (Figure S8). Effect sizes of LLD-AFR and CpG-content increased, due to opposing effects with the recombination rate and background selection statistic annotations. Effect sizes of the recombination rate, nucleotide diversity and background selection statistic annotations decreased because they compete with each other, and LLD and GC-content were no longer significant (after correction for multiple testing) due to their high correlation with LLD-AFR and CpG-content, respectively (Table S10). Psychiatric diseases and autoimmune diseases exhibited significantly stronger effects for the predicted allele age and background selection statistic annotations, respectively (Table S11), possibly due to the role of selection at different time scales in shaping the genetic architecture of these diseases^30,31^.

To provide a more intuitive interpretation of the magnitude of the LD-related annotation effects, we computed the proportion of heritability explained by each quintile of each annotation in the baseline-LD model, and by each quintile of MAF for comparison purposes (Figure 4, Table S9, Table S12, and Figure S9). These proportions are computed based on a joint fit of the baseline-LD model, but measure the heritability explained by each quintile of each annotation while including the effects of other annotations—in contrast to standardized effect sizes *τ*\*, which are conditioned on all other annotations and measure the additional contribution of one annotation to the model. The youngest 20% of common SNPs (based on MAF-adjusted predicted allele age) explained 3.9x more heritability than the oldest 20%. This is even larger than MAF-dependent effects, in which the 20% of common SNPs with largest MAF (> 38%) explain 1.8x more heritability than the 20% with smallest MAF (< 10%). (We note that slightly smaller heritability explained for less common variants is consistent with larger *per-allele* effect sizes for less common variants, as less common variants with the same per-allele effect size explain less heritability in proportion to *p*(1–*p*); see Discussion for additional comments on MAF-dependent effects.) The heritability explained by quintiles of recombination rate was roughly flat (in contrast to *τ*\*, which conditions on effects of other annotations; Figure 3c) due to the inclusion of opposing effects of the LLD-AFR and CpG-content annotations (Table S13 and Table S14); we note that the effect of recombination rate is dominated by its largest (5^th^) quintile (i.e. recombination rate hotspots, Figure S10), explaining the significant decrease in heritability explained between the 4^th^ and 5^th^ quintiles (Figure 4).

**Figure 4:**
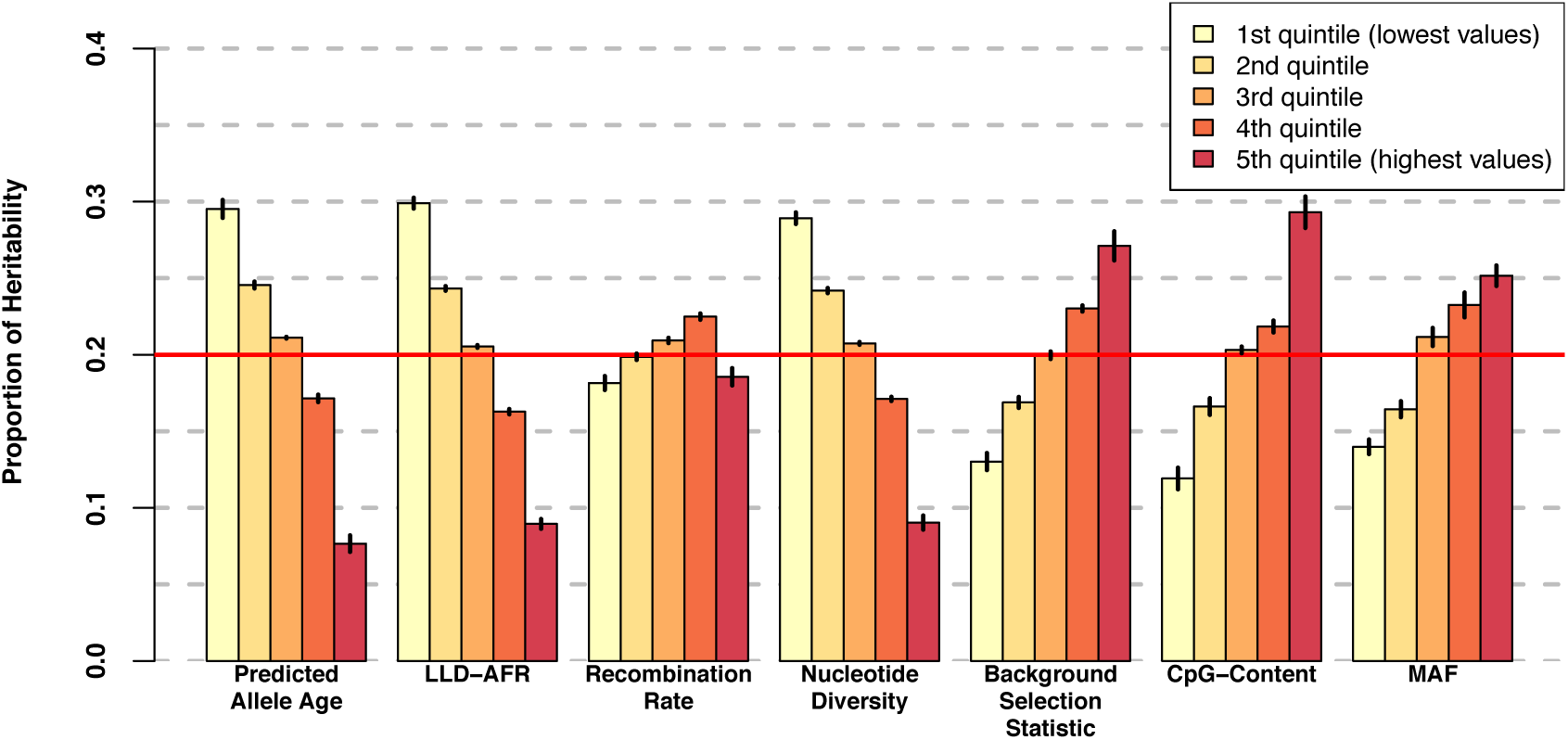
Proportion of heritability explained by the quintiles of each LD-related annotation, meta-analyzed over 31 independent traits. We report results for each LD-related annotation of the baseline-LD model, and for MAF for comparison purposes. Numerical results are reported in Table S12. Results for all 56 complex traits are reported in Figure S9 and Table S9. Error bars represent jackknife standard errors around the enrichment estimates. The red line indicates the proportion of heritability when there is no enrichment (20% of SNPs explain 20% of heritability).

### LD-related annotations predict deleterious effects

Our finding that common variants that are more recent tend to explain more complex trait heritability is potentially consistent with the action of negative selection on variants affecting complex traits, since selection has had less time to eliminate recent weakly deleterious variants. We hypothesized that our results for other LD-related annotations might also be explained by the action of negative selection. To investigate this hypothesis, we performed forward simulations^32^ using a demographic model for African and European populations^27^ and a range of selection coefficients for deleterious variants (see Online Methods). We jointly regressed the absolute value of the selection coefficient against the allele age (now using true allele age instead of predicted allele age), LLD-AFR, recombination rate and nucleotide diversity annotations from the baseline-LD model to assess whether these annotations are jointly predictive of deleterious effects (the background selection statistic and CpG-content annotations could not be investigated as they rely on empirical data). We observed that these four annotations were all significant in the joint analysis (Figure 5 and Table S15), with effect sizes roughly proportional to the standardized effect sizes for trait heritability reported in Figure 3c. This suggests that the joint impact of each of these annotations on trait heritability is a consequence of their predictive value for deleterious effects. Indeed, consistent with theory, recent variants are more likely to be deleterious since selection has had less time to remove them^33^, variants in low recombination rate regions are more likely to be deleterious due to reduced efficiency of selection (Hill-Robertson effect^28^), and variants in low nucleotide diversity regions are more likely to be deleterious due to increased efficiency of selection in those regions^34^. In addition, the LLD-AFR annotation contains information complementary to allele age, recombination rate and nucleotide diversity; we note that LLD-AFR contains roughly the same amount of information (i.e. the same effect) as LLD measured in an ancestral population sampled just before the out-of-Africa event (Figure S11). We further determined that the predictive value of the nucleotide diversity annotation is contingent on the non-homogeneous distribution of selection coefficients, and that the predictive value of the LLD-AFR annotation is largely contingent on the out-of-Africa bottleneck, as the LLD effect disappears in a constant population size model with a homogeneous distribution of selection coefficients (Figure S12). We finally note that we did not expect our results for LD-related annotations to be a signature of positive selection on variants affecting complex traits, as beneficial alleles tend to have increased LD^35^ and more efficient selection in high recombination rate regions^28^, each of which would be inconsistent with the results in Figure 3c; indeed, forward simulations involving beneficial mutations confirmed that the LD-related annotations associated to per-SNP heritability do not predict beneficial effects (Figure S13).

**Figure 5:**
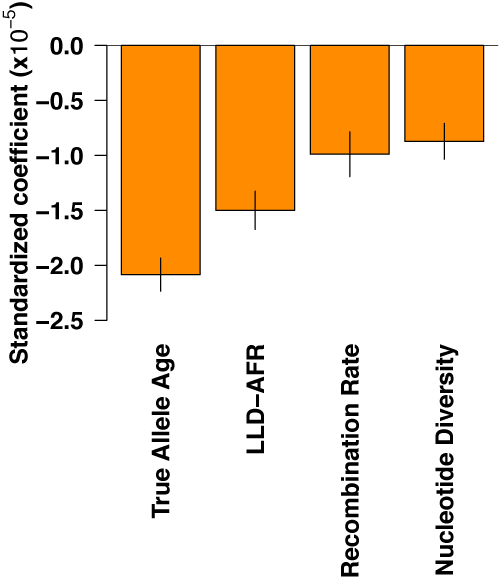
Forward simulations confirm that LD-related annotations predict deleterious effects. We report standardized coefficients for each of four LD-related annotations in a joint regression of absolute selection coefficient against these annotations in data from forward simulations (see text). Numerical results are reported in Table S15. Error bars represent 95% confidence intervals around the regression coefficient estimates.

## Discussion

In this study, we assessed the LD-dependent architecture of human complex traits by extending stratified LD score regression^8^ from binary to continuous annotations, an approach that produces robust results in simulations. We determined that SNPs with low LLD have larger per-SNP heritability across all 56 complex traits analyzed. More than half of this signal can be explained by functional annotations that are negatively correlated with LLD and enriched for heritability, such as DHS and histone marks. The remaining signal is largely driven by MAF-adjusted predicted allele age, as more recent alleles have larger per-SNP heritability in 55 out of the 56 complex traits analyzed, but we also observed multiple jointly significant effects of other LD-related annotations. We showed via forward simulations that all of these jointly significant effects are consistent with the action of negative selection on deleterious variants. As noted above, recent variants are more likely to be deleterious since selection has had less time to remove them^33^, variants in low recombination rate regions are more likely to be deleterious due to reduced efficiency of negative selection (Hill-Robertson effect^28^), and variants in low nucleotide diversity regions are more likely to be deleterious due to increased efficiency of selection in those regions^34^; we also observed higher per-SNP heritability for SNPs with low values of LLD-AFR, capturing a property of variant history that is currently unknown. We note that our genome-wide results on recombination rate differ from the results of Hussin et al.^16^, who determined that regions of low recombination rate are enriched for exonic deleterious and disease-associated variants: although we do observe a similar recombination rate effect (consistent with the Hill-Robertson effect^28^) for jointly estimated effect sizes *τ*\*, which are conditioned on other annotations and measure the additional contribution of one annotation to the model, this effect is largely canceled out when including the opposing effects of other annotations (Figure 4, Table S13 and Table S14).

While negative selection has long been hypothesized to shape genetic diversity^24^,and previous studies have emphasized the importance of allele age^21,36,37,^ and recombination rate^16^, our study demonstrates the impact of negative selection on complex traits on a polygenic genome-wide scale. Specifically, our results demonstrate that common variants associated to complex traits are weakly deleterious, confirming a hypothesized relationship between the effect size of a variant and its selection coefficient *s* (ref. ^38–41^). One of the implications of this finding is that we expect larger per-allele effect sizes for less common variants, consistent with only slightly smaller (per-SNP) heritability explained (Figure 4); this expectation also applies to rare variants, which we do not analyze here. We note that although we have focused here on unsigned heritability enrichment analyses, weakly deleterious derived alleles might systematically increase disease risk; we caution that signed analyses to assess this may be susceptible to confounding due to population stratification, as differences in demography may lead to systematic differences in derived allele frequencies across subpopulations.

Our results on LD-dependent architectures have several implications for downstream analyses. First, recent work has suggested that the problem of LD-related bias in SNP-heritability estimates^11,12^ could be addressed by modeling regional LD (LD-REG) in addition to MAF^13^. On the other hand, our baseline-LD model contains a considerably larger number of parameters, increasing model complexity but more accurately resolving the underlying signal; in particular, our results suggest that modeling predicted allele age may be more informative than modeling regional LD (Figure 4). Second, previous studies have shown limited improvements in polygenic prediction accuracy^7,42^ and association power^43,44^ using functional annotations, perhaps because the annotations analyzed in those studies have pervasive LD between in-annotation and out-of-annotation SNPs^7^; however, our LD-related annotations by definition should not have this limitation, making them potentially more useful in those contexts. Third, although SNPs with low LLD have larger causal effect sizes, SNPs with high LLD may have larger *χ^2^* statistics if they tag multiple causal variants. In the presence of multiple causal variants, fine-mapping strategies based on ranking *P* values^45^ might thus favor high-LLD non-causal variants over causal low-LLD variants. For this reason, approaches that explicitly model multiple causal variants while incorporating LD-dependent architectures using integrative methods^46^ might improve fine-mapping accuracy. Fourth, we observed that predicted allele age is substantially smaller (>0.1 standard deviations below average) in transcription start site (TSS), coding, conserved and UTR regions and below average for all functional annotations except repressed regions (Table S16), consistent with stronger selection. The identification of functional non-coding regions under strong selective constraint could be used to improve variant prioritization in whole-genome sequencing studies ^40,47^.

Although our work has provided insights on the genetic architecture of human complex traits, it has several limitations. First, our extension of stratified LD score regression assumes a linear effect of each continuous annotation (Equations (1) and (3), see Online Methods), which may not always hold; however, this assumption appears reasonable in the continuous annotations that we analyzed (Figure 4). Second, we restricted all of our analyses to common variants (see Online Methods), as stratified LD score regression has several limitations when applied to rare variants^8^. Third, as noted above, results of stratified LD score regression may be biased in the presence of unmodeled functional annotations^8^; we believe it is unlikely that this impacts our main conclusions, both because we included a large number of baseline model annotations in our analyses and because our results (Figure 3c) are consistent with selection effects in forward simulations (Figure 5). Fourth, while the allele age predictions produced by ARGweaver^21^ were of critical value to this study, they have > 10% missing data, were computed on only 54 sequenced individuals (including only 13 Europeans), and rely on a demographic model with constant population size; the development of computationally tractable methods for predicting allele age remains a research direction of high interest. Fifth, while the effect directions of the LD-related annotations we analyzed were remarkably consistent across all 56 complex traits analyzed, this result does not imply that negative selection acts directly on each of these traits, as selection may be acting on pleiotropic traits^48^. Sixth, while our results suggest that negative selection has a greater impact than positive selection on the genetic architecture of human complex traits, we cannot draw broader conclusions about the roles of negative and positive selection in shaping the human genome^49,50^. In addition, our forward simulations did not include balancing selection, whose main genomic signature (in contrast to negative selection) is increased nucleotide diversity^51^ and would not explain the results in Figure 3c, or stabilizing selection, which uses negative selection to favor intermediate values of phenotypes over extreme values. Seventh, the interpretation of some LD-related annotations remains unclear. The LLD-AFR annotation captures a property of variant history that is currently unknown. The CpG-content annotation is highly correlated to the GoNL local mutation rate map annotation^52^ (*r* = 0.86, Table S17), but that annotation does not have a significant effect on trait heritability when conditioned on the baseline model (Table S18), suggesting that the CpG-content annotation might instead tag some functional process absent from the baseline model; indeed, some of our LD-related annotations could be viewed as proxies for currently unknown functional annotations. Despite all of these limitations, our results convincingly demonstrate the action of negative selection on deleterious variants that affect complex traits, complementing efforts to learn about negative selection by analyzing much smaller rare variant data sets.

## Acknowledgements

We thank the research participants and employees of 23andMe for making this work possible. We thank S. Sunyaev, Y. Reshef, G. Kichaev, D. Speed and F. Day for helpful discussions. This research has been conducted using the UK Biobank Resource (Application Number: 16549). This research was funded by NIH grants R01 MH101244, R01 MH107649 and U01 HG009088.

## Online Methods

### Extension of stratified LD score regression to continuous annotations

The derivation of stratified LD score regression using binary annotations has previously been described^8^. Here, we extend the method to continuous-valued annotations.

Suppose that we have a sample of *N* individuals, and a vector *y* = (*y*_1_, …, *y_N_*) of quantitative phenotypes, standardized to mean 0 and variance 1. We assume the infinitesimal linear model

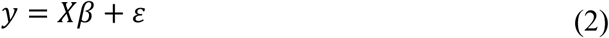
where *X* is a *N* × *M* matrix of standardized genotypes, *β* = (*β*_1_, …, *β_M_*) is the vector of per normalized genotype effect size, and *ε* = (*ε*_1_, …, *ε_N_*) is a mean-0 vector of residuals with variance 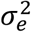. Here, we are interested in modeling *β* as a mean-0 vector whose variance depends on *C* continuous-valued annotations *a*_1_,…, *a_c_*:

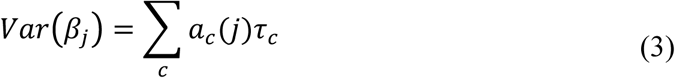
where *a_c_* (*j*) is the value of annotation *a_c_* at SNP *j*, and τ*_c_* represents the per-SNP contribution of one unit of the annotation *a_c_* to heritability. This is a generalization of stratified LD score regression^8^, with *a_c_*(*j*) ∈ {0,1} if annotation *a_c_* has binary values.

Let 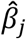 be the estimate of the marginal effect of SNP *j* in our sample. According to Finucane et al.^8^, we can write

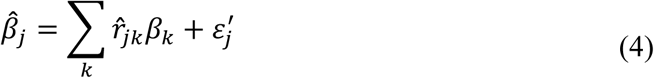
Where 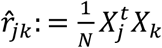 is the in-sample correlation between SNPs *j* and *k*, and 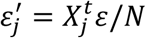 (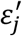 has mean 0 and variance 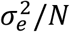).

We now consider the expectation of 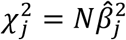. We can write 
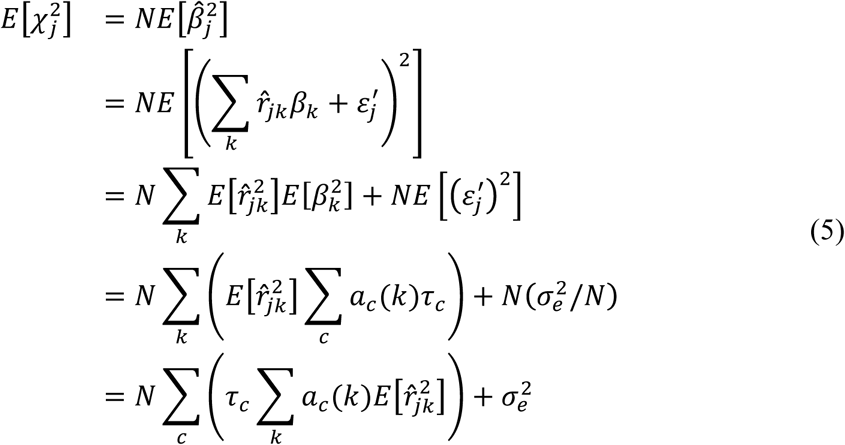
where the third equality holds because 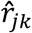, *β_k_*, and 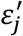 are independent and *β* and *ε′* have mean 0. Note that *r_jk_* denotes the true correlation between SNPs *j* and *k* in the underlying population and that *r_jk_* is fixed throughout, so that 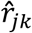 and *β_k_* are independent even though both depend on *r_jk_*. In an unstructured sample, we have 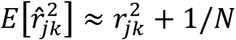. We thus have 
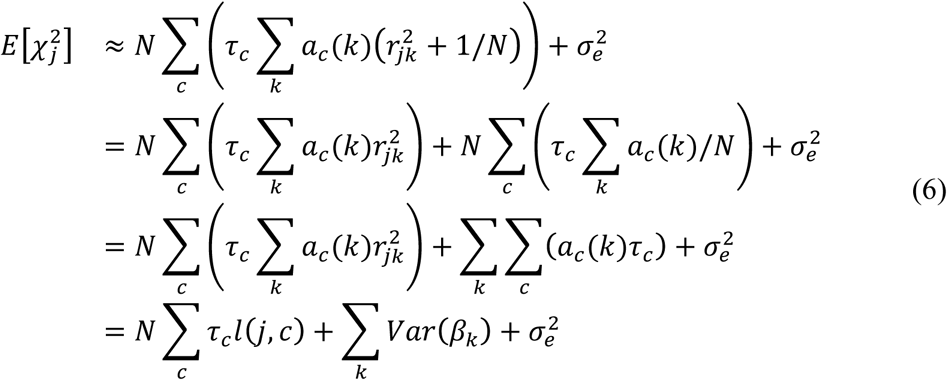
Where 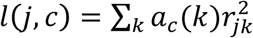 is the LD score of SNP *j* with respect to annotations *a_c_*. As the variance of our phenotype *y* is 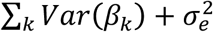 and is equal to 1 by definition, this reproduces the main equation of stratified LD score regression (modulo the term *Nb* for confounding biases): 
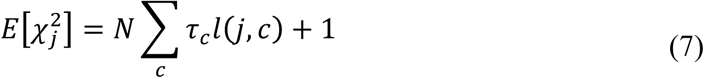
We were interested in both comparing the estimated effect size of the different annotations and meta-analyzing them across different traits. For this reason, we focused on per-standardized annotation effect sizes 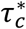, defined as the additive change in per-SNP heritability associated to a 1 standard deviation increase in the value of the annotation, divided by the average per-SNP heritability over all SNPs for the trait, and computed as

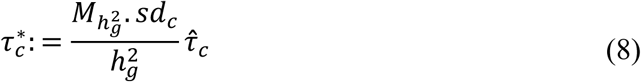
Where 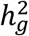 the estimated SNP-heritability of the trait computed as 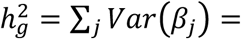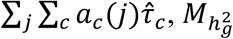 is the number of SNPs used to compute 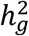 and *sd_c_* is the standard deviation of the annotation *a_c_*.

To interpret the heritability explained by a continuous-valued annotation *a_c_*, we computed the expected heritability of each quintile of its annotations. Let *C_c,q_* denote the *q*-th quintile of annotation *a_c_*, so that 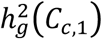 and 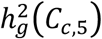 represent the heritability explained by the 20% of SNPs with the lowest and highest values of *C_c_*, respectively. We used the equation 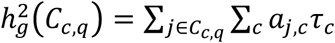 to estimate 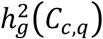.

Application of stratified LD score regression was performed using Finucane et al.^8^ guidelines and was restricted to data sets of European ancestry. Reference SNPs, used to estimate LD scores, were defined as the set of 9,997,231 biallelic SNPs with minor allele count greater or equal than five in the set of 489 unrelated and outbred European samples^53^ from phase 3 of 1000 Genomes Project (1000G)^17^ (see URLs). Regression SNPs, used to estimate the vector of *τ* from GWAS summary statistics, were defined as the set of 1,217,312 HapMap Project Phase 3 SNPs, used here as a proxy for well-imputed SNPs. SNPs with unusual *χ^2^* association statistics (larger than 80 or 0.0001*N*), as well as SNPs in the major histocompatibility complex (MHC) region (chr6:25Mb-34Mb) were removed from all analyses. We note that the choice of regression SNPs is distinct from the choice of reference SNPs, and that regression SNPs tag potentially causal reference SNPs via LD scores computed using reference SNPs (see ref. ^8^ for further details). Heritability SNPs, used to compute *sd_c_*, 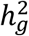 and 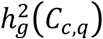, were the set of 5,961,159 reference SNPs with MAF ≥ 0.05. To assess the reproducibility of our results, we also considered 3,567 individuals of UK10K database^29^ (ALSPAC and TWINSUK cohorts) as a reference panel. We had 13,326,465 reference SNPs and 5,353,593 heritability SNPs in this analysis.

### Baseline model and functional annotations

The 59 functional annotations that we used to define the baseline model consist of the 53 binary annotations from ref. ^8^ and an additional six annotations. The 53 annotations are derived from 24 main annotations including coding, UTR, promoter and intronic regions, the histone marks monomethylation (H3K4me1) and trimethylation (H3K4me3) of histone H3 at lysine 4, acetylation of histone H3 at lysine 9 (H3K9ac) and two versions of acetylation of histone H3 at lysine 27 (H3K27ac), open chromatin as reflected by DNase I hypersensitivity sites (DHSs), combined chromHMM and Segway predictions (which make use of many Encyclopedia of DNA Elements (ENCODE) annotations to produce a single partition of the genome into seven underlying chromatin states), regions that are conserved in mammals, super-enhancers, and FANTOM5 enhancers. The 53 annotations also include 500bp windows around each of the 24 main annotations, 100bp windows around four of the main annotations, and an annotation containing all SNPs. We added four binary annotations based on super enhancers and typical enhancers^54^, as previously described^19^. We also added two conserved annotations based on GERP++ scores^55^, including one continuous annotation based on the neutral rate (NS) score and one binary annotation based on a rejected substitutions (RS) score ≥ 4, as we observed significant effects for these annotations (see Table S19). We did not include 500bp windows around the GERP-NS annotation (which is a continuous annotation) or the GERP-RS annotation (which is defined separately for each base pair).

### MAF adjustment and LLD annotations

To investigate the LD-dependent architecture of human complex traits, it is essential to account for the relationship between minor allele frequency (MAF) and LD. Indeed, common variants have both higher LD scores and per-SNP heritability^5,9^. For this reason, all of our stratified LD score regression analyses included 10 MAF bins coded as 10 binary annotations (all with MAF ≥ 0.05, see Table S20) in addition to an annotation containing all SNPs.

To quantify the level of LD (LLD) of reference SNPs, we first computed LD scores, defined as the sum of squared correlations of each SNP with all nearby SNPs in a 1 cM window, using the ldsc software. Then, we MAF-adjusted these values via MAF-stratified quantile normalization: for each MAF bin, LD scores were quantile normalized to a normal distribution of mean 0 and variance 1. The LLD of rare variants (MAF < 0.05) was fixed to 0. Because stratified LD score regression is designed to quantify the heritability explained by common SNPs, and the heritability explained by rare variants is hypothesized to be relatively low^1,5,56^, we excluded rare variants from all MAF-adjusted annotations. For the LLD model (Figure 1), we thus modeled the variance of the per-normalized genotype effect size of SNP *j* as:

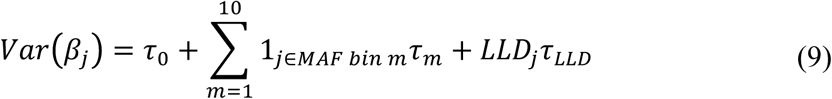
where *τ*_0_ is an intercept term modeling the per-SNP contribution of each SNP to heritability, 1_*j∈MAF bin m*_ is an indicator function with value 1 if SNP *j* belongs to MAF bin *m* and 0 otherwise, *τ_m_* is the per-SNP contribution of a SNP in MAF bin *m* to heritability, and *τ_LLD_* is the contribution of one unit of the annotation LLD to heritability.

The LLD-D’ annotation of a SNP was measured by summing the D’ coefficients of that SNP with all nearby SNPs in a ±0.5 cM window. Version 1.90b3 of PLINK 2 software^57^ (see URLs) was used to compute D’ coefficients for each pair of SNPs. The LLD of a genomic region (LLD-REG) was measured by averaging in 100 kb windows the LD scores computed in 20-Mb regions (ignoring LD *r*^2^ < 0.01), as previously described^13^, using the ‐‐ld-score-region option of version 1.25.1 of GCTA software^2^. LLD-D’ was MAF-adjusted via MAF-stratified quantile normalization. LLD-REG was quantile normalized without MAF-adjustment because it is a regional annotation.

The LLD-AFR annotation was measured by computing LD scores of reference SNPs in 440 unrelated African samples from phase 3 of 1000 Genomes Project (ACB and ASW populations were removed due to the presence of European admixture). LD scores for reference SNPs that were absent in African samples were set to 1. LLD-AFR was also MAF-adjusted via MAF-stratified quantile normalization, using the same European MAF bins.

### Other LD-related annotations

We used allele age as predicted by the ARGweaver^21^ method, estimated using 54 unrelated sequenced individuals (including 13 Europeans; see URLs). This annotation was also MAF-adjusted via MAF-stratified quantile normalization, as common variants tend to be older (the correlation for common reference SNPs between available ARGweaver allele ages and MAF is 0.16). 10.2% of common reference SNPs had missing values for predicted allele age; these values were excluded during the MAF-stratified quantile normalization process, and corresponding MAF-adjusted predicted allele ages were set to 0. Adding a binary annotation indicating missing allele age information for common reference SNPs did not change the effect size estimates for predicted allele age (Table S21).

Recombination rates, diversity, GC-content and CpG-content were computed using windows of different sizes: ± 10kb, ± 50kb, ± 100kb, ± 500kb, and ±1,000 kb. Recombination rates (measured in cM/Mb) were computed from three recombination maps (see URLs): the Oxford map, which estimates recombination rates from LD patterns in African, European and Asian populations from HapMap2^22,23^; the African-American map, which estimates recombination rates from admixture patterns in African-American individuals^58^; and the deCODE map, which estimates recombination rates from Icelandic parent-offspring pairs^59^. These recombination maps measure recombination rates at different time scale: the deCode map measures recombination that occurred in recent generations, the African-American map measures recombination that occurred in the past ∼20 generations, and the Oxford map measures recombination that occurred further back in time. The genetic positions of surrounding windows were interpolated linearly from recombination maps using PLINK. We determined that the Oxford map provided the most significant results (Table S3), suggesting that the impact of recombination rate on trait heritability operates over a long time scale; we thus used the Oxford map in all primary analyses. Nucleotide diversity was measured as the number of reference SNPs (with minor allele count ≥5) per kilobase. Measuring diversity on all 1000G SNPs (down to singletons or doubletons) or the fraction of rare variants^60^ (i.e. diversity of rare variants with allele count < 5) did not furnish more significant results (data not shown). GC-content and CpG-content were measured using version 2.17.0 of bedtools software^61^ and the human reference sequence used for the 1000 Genomes project (see URLs).

The background selection statistic was computed as 1 - McVicker B statistic^24^ to facilitate the interpretation of the results. Background selection statistic values close to 1 represent near complete removal of diversity as a result of background selection, and values near 0 indicate little effect. Replication timing was based on the Koren et al. annotation^25^. 0.19% and 0.27% of reference SNPs had missing values for background selection and replication timing, respectively; these were replaced by the median annotation value based on the remaining reference SNPs.

Finally, telomeres and centromeres were defined using window sizes of 5, 10 and 15 Mb, as described by Smith et al.^15^.

We thus created 43 LD-related annotations in total (see Table S22). For annotations computed with different windows sizes or using different data sources for recombination rate, the one producing the most significant *P* value after conditioning on the baseline model was selected as the primary annotation (Table S3). Except telomere and centromere annotations that were not significant in this analysis, other annotations had consistent results with adjacent window sizes. To overcome over-fitting, we used a Bonferroni threshold of 0.05 / 43 = 1.16 × 10^−3^ to assess statistical significance when analyzing one LD-related annotation at a time. We note that this procedure did not affect our final conclusions (see Figure S5).

### Choice of traits for main analyses and meta-analysis

Stratified LD score regression was applied to 29 publicly available GWAS summary statistic data sets^62-82^ (for age at menopause^80^, effect sizes are publicly available but sample sizes for each SNP were obtained through collaboration), 18 summary statistic data sets from 23andMe, and summary statistic of 15 traits from UK Biobank (see below). This led to total of 62 summary statistic data sets spanning 56 traits (five traits were represented in multiple data sets) with an average sample size of 101,401 (computed using the largest single data set for each trait; the average sample size of the 62 data sets is 101,989). Analyses were restricted to traits for which the *z* score of total SNP-heritability computed using the baseline model was at least 6 (Table S1). Traits displayed in Figure 1 were selected by prioritizing them according to the total SNP-heritability, excluding traits with absolute genetic correlation > 0.50 (ref. ^20^). Traits included in the meta-analyses were selected by prioritizing them according to the *z* score of total SNP-heritability and excluding genetically correlated traits in overlapping samples by measuring the intercept of cross-trait LD score regression^20^ as previously described^8^. We retained 31 independent traits (average *N* = 84,686, Table S1) and performed random-effects meta-analyses using the R package *rmeta*.

For analyses of psychiatric and autoimmune diseases, we considered five psychiatric diseases with low sample overlap (anorexia, autism, bipolar disorder, depressive symptoms and schizophrenia) and six autoimmune diseases with low sample overlap (celiac, cirrhosis, eczema, lupus, inflammatory bowel disease and rheumatoid arthritis). We meta-analyzed standardized effect sizes *τ\** for the five psychiatric diseases and six autoimmune diseases using random effects, and compared the results with results for non-psychiatric and non-autoimmune diseases using a *t*-test. Non-psychiatric and non-autoimmune diseases were defined by removing psychiatric diseases and autoimmune diseases from the set of 31 independent traits, leading to a total of 28 and 29 traits, respectively.

### 23andMe data set

For the 23andMe study, participants were drawn from the customer base of 23andMe Inc. (Mountain View, CA), a consumer genetics company^83,84^. All participants included in the analyses provided informed consent and answered surveys online according to the 23andMe human subjects protocol, which was reviewed and approved by Ethical & Independent Review Services, a private institutional review board. Samples were genotyped on one of four genotyping platforms. The V1 and V2 platforms were variants of the Illumina HumanHap550+ BeadChip, including about 25,000 custom SNPs selected by 23andMe, with a total of about 560,000 SNPs. The V3 platform was based on the Illumina OmniExpress+ BeadChip, with custom content to improve the overlap with our V2 array, with a total of about 950,000 SNPs. The V4 platform in current use is a fully custom array, including a lower redundancy subset of V2 and V3 SNPs with additional coverage of lower-frequency coding variation, and about 570,000 SNPs.

Participants were restricted to a set of individuals who have > 97% European ancestry, as determined through an analysis of local ancestry^85^. A maximal set of unrelated individuals was chosen for each analysis using a segmental identity-by-descent (IBD) estimation algorithm^86^. Individuals were defined as related if they shared more than 700 cM IBD, including regions where the two individuals share either one or both genomic segments identical-by-descent. This level of relatedness (roughly 20% of the genome) corresponds approximately to the minimal expected sharing between first cousins in an outbred population.

Participant genotype data were imputed against the March 2012 “v3” release of 1000 Genomes reference haplotypes, phased with ShapeIt2 (ref. ^87^). Data were phased and imputed for each genotyping platform separately. Data were phased using a 23andMe developed phasing tool, Finch, which implements the Beagle haplotype graph-based phasing algorithm^88^, modified to separate the haplotype graph construction and phasing steps.

In preparation for imputation, phased chromosomes were split into segments of no more than 10,000 genotyped SNPs, with overlaps of 200 SNPs. SNPs with Hardy-Weinberg equilibrium *P* < 10^−20^, call rate < 95%, or with large allele frequency discrepancies compared to European 1000 Genomes reference data were excluded. Frequency discrepancies were identified by computing a 2x2 table of allele counts for European 1000 Genomes samples and 2000 randomly sampled 23andMe participants with European ancestry, and identifying SNPs with a chi squared *P* < 10^−15^. Each phased segment was imputed against all-ethnicity 1000 Genomes haplotypes (excluding monomorphic and singleton sites) using Minimac2 (ref. ^89^), using 5 rounds and 200 states for parameter estimation.

The genetic association tests were performed using either linear or logistic regression as required assuming an additive model for allelic effects and controlled for age, sex, and five principal components of genetic ancestry.

### UK Biobank data set

We analyzed data from the UK Biobank (see URLs) consisting of 152,249 samples genotyped on ∼800,000 SNPs and imputed to ∼73 million SNPs. One individual who had withdrawn consent was removed, leaving 152,248 samples (see URLs, Genotyping and QC). We selected 15 phenotypes with large sample size. For each phenotype, we computed mixed model association statistics on up to 145,416 European-ancestry samples using version 2.2 of BOLT-LMM software^90^ (see URLs) with genotyping array (UK BiLEVE / UK Biobank) and assessment center as covariates. We included 607,518 directly genotyped SNPs in the mixed model (specifically, all autosomal biallelic SNPs with missingness < 2 % and consistent allele frequencies between the UK BiLEVE array and the UK Biobank arrays), and we computed association statistics on imputed SNPs in HapMap3 (1,186,683 SNPs on average over the 15 phenotypes). Heritability enrichment analyses of UK Biobank data were based on analyses of summary statistics, despite the availability of individual-level data, both to ensure consistency with the remaining 48 summary statistic data sets and because we are not currently aware of a heritability enrichment method applicable to individual-level data that can analyze a large number of overlapping or continuous-valued annotations.

### Construction of the baseline-LD model

We first considered a model including the eight LD-related annotations that were significant after being conditioned on 10 MAF bins and the baseline model (i.e. LLD, predicted allele age, LLD-AFR, recombination rate, nucleotide diversity, the background selection statistic, GC-content and CpG-content), and also including 10 MAF bins and the baseline model. We removed LD-related annotations that were not significant (in the meta-analysis of 31 independent traits) one at a time based on the least significant *P* value (GC-content was first removed, then LLD). This procedure produced a baseline-LD model with the 59 annotations of the baseline model, the 10 MAF bins, and 6 remaining LD-related annotations, leading to a total of 75 annotations. We have made these annotations publicly available (see URLs).

### Simulations to assess extension of stratified LD score regression to continuous LD-related annotations

To ensure that applying our extension of stratified LD score regression to continuous LD-related annotations does not produce false-positive signals or biased results, we simulated quantitative phenotypes from chromosome 1 UK10K data^29^ (3,567 individuals and 1,041,378 SNPs). In each simulation, we used 1000G as the reference panel, and evaluated all 6 LD-related annotations of the baseline-LD model (Figure 3c), as well as the LLD annotation. We also included an annotation containing all SNPs and annotations for 10 MAF bins. In each simulation, we set trait heritability to *h*^2^ = 0.5 and selected *M* = 100,000 causal SNPs. Causal SNPs were selected randomly rom the 673,779 SNPs present in both UK10K and 1000G, such that all causal SNPs were represented in the reference panel. In null simulations, the variance of per-normalized genotype effect sizes was set to 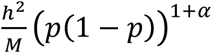 for a variant of frequency *p*. We considered simulations with both MAF-independent (*α* = -1, i.e. all SNPs have the same contribution to variance) and MAF-dependent architectures (*α* = -0.28, as previously estimated^9^). In causal simulations (MAF+LD-dependent architecture), we used the *τ* coefficients estimated from the meta-analyses reported in Figure 3a and set the variance of per-normalized genotype effect sizes using the additive model of equation (9), replacing the LLD annotation with the LD–related annotation of interest. These coefficients were rescaled to constrain the variance of each SNP to be positive and the total *h^2^* of the 100,000 causal SNPs to be 0.5. Phenotypes were simulated with GCTA^2^ (see URLs). 10,000 simulations were performed for each of the three simulation scenarios (null MAF-independent, null MAF-dependent, and causal MAF+LD-dependent). In each simulation, we estimated the effect size 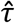 using HapMap 3 SNPs as regression SNPs, to account for the possibility that causal SNPs are not included in the set of regression SNPs. Corresponding 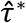, were computed using the simulated *h*^2^. (We were interested in the bias of the *τ* parameter, and not in the 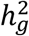 parameter which might be underestimated in simulation scenarios where rare variants have large effect sizes. We note that estimates of **τ\*** in real phenotypes may be slightly biased by inaccurate 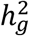 estimates, but that this will not lead to false-positive nonzero *τ* * estimates). We observed unbiased estimates of **τ\*** for most annotations in both null simulations (Figure 6a and Figure S14) and causal simulations (Figure 6b) (numerical results in Table S23). Only the recombination rate annotation exhibited very slight biases (between -0.028 and -0.025) that are nevertheless far from the estimates observed on real data (-0.540; Figure 3a). We also confirmed accurate calibration of standard errors in both null and causal simulations (Table S24). We repeated each of these simulations drawing causal SNPs from all UK10K SNPs (to simulate a scenario where causal SNPs are not represented in the reference panel). Results for null simulations were similar to above, and results for causal simulations produced slight biases opposite to (i.e. slightly underestimating) true effects (Figure S15; numerical results in Table S25).

**Figure 6:**
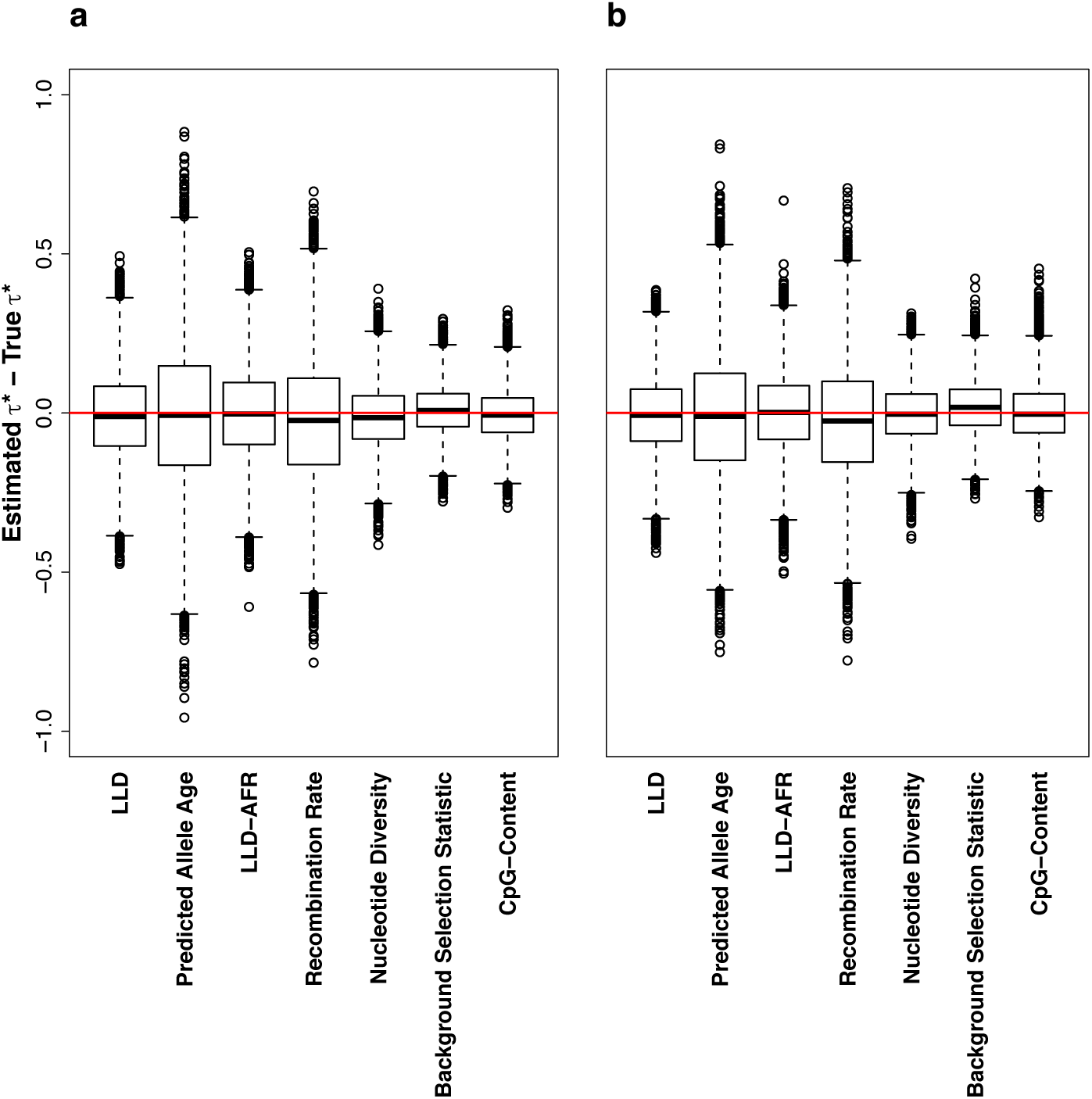
Simulations to assess extension of stratified LD score regression to continuous LD-related annotations. We report bias (estimated vs. true *τ\**) across 10,000 simulations for (a) Null simulations with MAF-dependent architecture and (b) Causal simulations with MAF+LD-dependent architecture. Results for null simulations with MAF-independent architecture are reported in Figure S14. Numerical results are reported in Table S23. Results for simulations with causal SNPs that are absent from the reference panel are reported in Figure S15 and Table S25.

### Coalescent simulations to assess the link between LLD and allele age

Coalescent simulations were performed using ARGON software^26^ (see URLs) to assess the correlation between the LLD and MAF-adjusted allele age of a SNP. We used demographical model parameters estimated in Gravel et al.^27^ to simulate European and African human genetic data, and assumed a generation time of 25 years. Recombination rate was set to 1 cM/Mb and mutation rate to 1.65 × 10^−8^ (ref. ^91^). We generated 33 fragments of 100 Mb for 500 European and 500 African individuals, representing a realistic genome size and sample sizes equivalent to the reference populations of 1000G. LD scores were computed independently in each 100 Mb fragment on SNPs with an allele count ≥ 5 in Europeans, and allele age and LD scores were MAF-adjusted via MAF-stratified quantile normalization after merging the 33 fragments.

### Forward simulations to assess the connection between LD-related annotations and negative selections

To investigate the connection between the LD-related annotations of the baseline-LD model (predicted allele age, LLD-AFR, recombination rate and nucleotide diversity; note that background selection statistic and CpG-content cannot be assessed in simulations as they rely on empirical data, and that these simulations used true allele age instead of predicted allele age) and the selection coefficient *s*, we performed forward simulations under a Wright-Fisher model with selection using version 1.8 of SLiM software^32^ (see URLs). We simulated 1Mb regions of genetic length 1cM. To ensure realistic recombination rate patterns, we divided the 1Mb regions into three recombination environments^16^, including a coldspot region of 475 kb containing 4.1% of recombination events (i.e. 0.08 cM/Mb) and a high recombination rate region of 140 kb containing 58.6% of recombination events (4.18 cM/Mb). The mutation rate was again set to 1.65 × 10^−8^ (ref. ^91^). New mutations had probability *d* to be deleterious with a dominance coefficient of 0.5 and a selection coefficient *s* drawn from a gamma distribution with parameters -0.05 and 0.2 (as suggested in the SLiM manual), and 1 – *d* to be neutral (i.e. *s* = 0). To study the impact of a non-homogeneous distribution of *d* across the genome, we divided each recombination environment into two sub-regions and assigned alternate probabilities *d_1_* and *d_2_* to be deleterious in these sub-regions (results reported in Figure 5 used *d_1_* = 0.60 and *d_2_* = 0.90). We performed simulations spanning 100,000 generations under 2 different demographic scenarios. First, we started from a fixed population size of 7,300 individuals, used the realistic demographic model of Gravel et al.^27^ for the last 5,920 generations, and outputted 500 European genomes and 500 African genomes. Second, we considered a fixed population size of 10,000 individuals and outputted 500 individual genomes at the last generation. We simulated 200 1Mb regions in each demographic scenario. LD scores were computed independently in each 1Mb fragment based on SNPs with a minor allele count ≥ 5; allele age and LLD-AFR or LLD (depending on the demographic scenario) were MAF-adjusted via MAF-stratified quantile normalization after merging the 200 1Mb regions. We performed a multivariate linear regression of the absolute value of the (known) selection coefficients |*s*| against the MAF bins annotations, the MAF-adjusted allele age, the MAF-adjusted LLD-AFR or LLD (depending on the demographic scenario), the true recombination rate, and the nucleotide diversity measured in a ± 10 kb window size. The above simulations did not include beneficial mutations, but we also performed simulations with beneficial and neutral mutations only to confirm that positive selection cannot explain the observed results of the LD-related annotations of the baseline-LD model (Figure S12).

## URLs

ldsc software, http://www.github.com/bulik/ldsc;

baseline-LD annotations, https://data.broadinstitute.org/alkesgroup/LDSCORE/;

1000 Genomes Project Phase 3 data, ftp://ftp.1000genomes.ebi.ac.uk/vol1/ftp/release/20130502;

PLINK software, https://www.cog-genomics.org/plink2;

ARGweaver allele ages, http://compgen.cshl.edu/ARGweaver/CG_results/download;

Oxford recombination map, http://www.shapeit.fr/files/genetic_map_b37.tar.gz;

African-American and deCode recombination maps,

http://www.well.ox.ac.uk/∼anjali/AAmap/maps_b37.tar.gz;

bedtools software, http://bedtools.readthedocs.org/en/latest;

Human reference sequence, ftp://ftp-trace.ncbi.nih.gov/1000genomes/ftp/technical/reference/human_g1k_v37.fasta.gz;

GCTA software, http://cnsgenomics.com/software/gcta/download.html;

ARGON software, https://github.com/pierpal/ARGON;

BOLT-LMM software, https://data.broadinstitute.org/alkesgroup/BOLT-LMM;

UK Biobank, http://www.ukbiobank.ac.uk/;

UK Biobank Genotyping and QC Documentation, http://www.ukbiobank.ac.uk/wp-content/uploads/2014/04/UKBiobank_genotyping_QC_documentation-web.pdf;

SLiM software, https://messerlab.org/slim/;

